# Fast Ripple-Delta Coupling as Early Biomarker for Post-Traumatic Epileptogenesis in Repetitive Brain Injury

**DOI:** 10.1101/2025.09.16.676387

**Authors:** Oleksii Shandra, Dzenis Mahmutovic, Biswajit Maharathi, Md Adil Arman, Michael J Benko, Owen Leitzel, Pritom Kumar Saha, Stefanie Robel

## Abstract

Traumatic brain injury (TBI) can induce post-traumatic epilepsy (PTE), but early biomarkers for epileptogenesis are lacking. We used a repetitive diffuse TBI (rdTBI) model in mice with continuous video-EEG monitoring up to 4½ months post-injury to investigate electrographic biomarkers before and during post-traumatic seizure development. 25% of mice developed post-traumatic seizures with highly variable latency (5-126 days post-injury). Most significantly, we identified fast ripple-delta DOWN state coupling as an early biomarker that was detectable at 4 days post-TBI and appeared before seizure onset in all seizure-experiencing mice. This EEG signature distinguished seizure-experiencing from seizure-free TBI mice with high specificity. Power spectrum analysis revealed elevated delta and theta power, reduced physiological fast oscillations (alpha, beta, gamma) and increased pathological high-frequency oscillations (fast ripples) in seizure-experiencing animals, indicating network hyperexcitability. Spike analysis showed that while TBI itself increased cortical excitability, seizure onset triggered a dramatic further escalation in interictal activity. These electrographic signatures were remarkably consistent across all seizure-experiencing animals regardless of single or recurrent seizure pattern. Our results demonstrate that fast ripple-delta coupling represents a promising early biomarker detectable at 4 days post-TBI, before seizure onset, offering potential for early identification of post-traumatic seizure susceptibility. Importantly, this biomarker identified all seizure-prone animals regardless of whether they developed single or recurrent seizures, suggesting shared underlying mechanisms and clinical relevance for any post-traumatic seizure occurrence. These findings emphasize the utility of temporal EEG analysis for detecting early electrographic changes in post-traumatic epileptogenesis and may inform future intervention strategies.

**Key Points:**

- Fast ripple-delta DOWN state coupling was detectable as early as 4 days post-TBI and appeared before seizure onset in seizure-experiencing mice, representing the first early biomarker that can stratify animals for epileptogenesis risk during the critical latent period.
- Delta and theta power increased while alpha, beta and gamma power decreased in all seizure-experiencing mice post-TBI, creating a consistent electrographic signature regardless of whether animals developed single or recurrent seizures.
- Fast ripples were elevated and gamma-to-HFO ratios were reduced in seizure-experiencing mice, reflecting network hyperexcitability shift and potential inhibitory dysfunction that preceded seizure onset.
- Seizure onset triggered a 3-fold escalation in spike activity, while baseline spike differences between TBI and pre-seizure mice were not significant, highlighting the limitation of spike counts alone as predictive biomarkers during the latent period.
- Electrographic signatures were almost similar across all seizure patterns (single and recurrent), suggesting shared underlying mechanisms of network dysfunction, though larger studies are needed to determine if biomarkers can predict seizure frequency in addition to seizure susceptibility.

## Introduction

Post-traumatic epilepsy (PTE) is a particularly debilitating chronic disorder induced by traumatic brain injury (TBI) characterized by unprovoked, recurrent seizures^1, 2^. Forty percent of patients who develop PTE experience their first late seizure within the first 6 months after TBI and 80% of PTE patients will have had their first late seizure within 2 years after injury^3^. Currently, it is impossible to determine early after TBI which patients will ultimately develop PTE.

Spontaneous seizures have been confirmed in rodent TBI models including fluid percussion injury (FPI)^4–6^, controlled cortical impact (CCI)^6–8^, weight drop^9, 10^ and blast injury^11–13^. These models recapitulate key pathologies including cell death^14^, blood-brain barrier breakdown^15^ and glial activation^7^. However, PTE incidence varies considerably across models, ranging from ∼3% in lateral FPI to 50-58% in CCI and weight drop models^6, 8–10, 16^.

Since not all animals that undergo TBI develop PTE, it is critical to distinguish animals that develop seizures from those that do not before identifying molecular and cellular mechanisms that cause PTE. We previously found that mice that developed PTE had larger areas of atypical astrocytes compared to mice that did not develop PTE^10^. Atypical astrocytes rapidly lose homeostatic proteins involved in maintaining neuronal function including the potassium channel Kir4.1 and the glutamate transporter Glt1, which may contribute to the development of PTE^10^. However, identifying PTE mice required months of EEG recordings post-TBI, with tissue changes examined after PTE onset, making it impossible to determine if detected changes cause or result from PTE. Establishing methods to identify PTE-susceptible mice earlier, ideally before seizure onset, is essential since repeated seizures drive additional neuroinflammation and pathological changes.

EEG forms the basis for epilepsy diagnosis and management^17^, with the latency period following TBI representing a critical window for early intervention before PTE onset. High-frequency oscillations (HFOs), particularly fast ripples (250-1000 Hz), have emerged as promising biomarkers of the epileptogenic brain^18–21^. Fast ripples represent pathological synchronization of cellular assemblies related to seizure onset zones^18^ and their synchronization with delta peaks may be predictive for epilepsy^18^ by reflecting coordinated neural activity preceding epileptic events. However, whether fast ripple-delta coupling can predict PTE in animal models remains unassessed.

Here, we used a model of repeated mild, diffuse TBI (rdTBI) caused by weight-drop^9^. We previously reported the development of PTE in a subset of mice in this model in the absence of primary tissue loss, pronounced glial activation and hemorrhage^9, 10^. The primary goal of the present study was to identify reliable electrographic biomarkers for early post-traumatic seizure detection. We aimed to identify EEG biomarker profiles that precede and predict seizure onset, with particular focus on fast ripples and their coupling with delta waves before seizure onset in TBI mice that developed post-traumatic seizures.

## Materials and Methods

### Experimental design and exclusion criteria

42 mice were randomly assigned to TBI or Sham groups. Continuous 24/7 recordings were conducted for up to 4.5 months post-TBI, though 9 mice (6 TBI, 3 sham) were monitored for 2 months starting at 2 months post-TBI. Exclusion criteria included electrode site hemorrhage, lost EEG caps, signal loss, or noise corrupting >50% of recordings. One sham mouse was excluded due to reference electrode hemorrhage.

### Animals

10- to 16-week-old male C57BL/6 mice (bred in-house or purchased from *Jackson Laboratory*) were used. All animal procedures were approved and performed according to the guidelines of the Institutional Animal Care and Use Committee of Virginia Polytechnic Institute and State University (*Virginia Tech*) and were conducted in compliance with the National Institutes of Health’s Guide for the Care and Use of Laboratory Animals.

### Weight-drop injury model

Mice received repetitive diffuse TBI based on an established protocol^9, 10^. Briefly, mice were anesthetized and placed on a foam pad, with a metal disc on their head. Buprenorphine was administered at a dose of 0.05-0.1 mg/kg subcutaneously before the procedure. TBI group mice received three impacts from a 50 cm height with 45-minute intervals between impacts. This injury modality induced mild/concussive injury as previously validated^10^. Sham mice underwent isoflurane anesthesia, administration of buprenorphine and placement in the weight drop device but not the impact. Naïve mice were excluded from these procedures.

### Stereotactic surgery

After one hour recovery period, mice were anesthetized with isoflurane and placed in a stereotactic apparatus for implantation of intracranial stainless steel screw electrodes with wire leads (00-96 x 1.6 mm, Plastics1) for either single-channel or multi-channel tethered recording. Depending on electrode configuration and data acquisition system configuration, monopolar (referential) or bipolar montage were used. A step-by-step protocol of the TBI and EEG implantation procedure was previously reported^9^.

### Video-EEG data acquisition

Mice were housed individually in monitoring cages. Two EEG systems were used: Pinnacle Technology and Biopac Systems. Continuous, 24/7 video-EEG data acquisition was performed from day one post-TBI for up to 19 weeks. In the Pinnacle Technology system, video-synchronized EEG data were collected at a sampling rate of 2000 Hz using Sirenia Acquisition software. In the *Biopac Systems*, video-synchronized EEG data were collected at a sampling rate of 500 Hz for single channel EEG and 2000 Hz for six-channel EEG using AcqKnowledge 4.1 software.

### Seizure detection and characteristics

EEG data and video analysis was conducted by an investigator blinded to the experimental groups. Electrographic seizures were defined using the following inclusion criteria: 1) duration of at least five seconds; 2) signal evolution in amplitude (at least three times compared to background activity) and frequency (3-12 Hz ictal spikes 20-70 milliseconds). These are separated by prolonged “interictal” periods during which the EEG trace is relatively stable. For each mouse, seizure incidence, frequency and duration were documented. All mice with at least one seizure, early or late, were included in the PTE group. To determine whether seizures were electroclinical or exclusively electrographic, a blinded investigator scored video traces corresponding to the electrographic seizures according to the modified Racine scale.^22^ EEG data underwent preprocessing including 60 Hz powerline interference removal using an infinite impulse response (IIR) filter, followed by bandpass filtering (2-10 Hz and 20-50 Hz) with 4th order IIR zero-phase digital filters. Data was segmented into 10-minute intervals for artifact detection using median and standard deviation thresholds. Shannon entropy was calculated on 0.5-second segments, with values exceeding one standard deviation from the median processed as potential seizures. Segments within 2.5 seconds were consolidated, followed by seizure edge detection using upper and lower envelope calculations. Final validation required signal power at least 50% higher than surrounding 10-minute periods. Reviewers blinded to experimental conditions cross-verified all seizure annotations.

### Spike Detection

Interictal spikes were detected and quantified using NeuroScore software with dynamic thresholding method. EEG data were preprocessed by applying a 60 Hz notch filter to remove powerline interference, followed by bandpass filtering (3-70 Hz). Automated spike detection was performed using dynamic thresholding parameters (minimum 4× baseline RMS; maximum 15× RMS), with an minimum value of 100 µV and spike width 30–70 ms)^23, 24^ optimized for interictal epileptiform discharges. All automatically detected spike events were subsequently verified through manual visual inspection by a trained investigator to ensure accuracy and eliminate false positives.

### Power Spectrum Analysis

The average spectral power within specific frequency bands, including a wide range (1-500 Hz), delta (1-4 Hz), theta (4-8 Hz), alpha (8-12 Hz), beta (12-30 Hz), gamma (30-50 Hz), high frequency oscillation (HFO, 80-250 Hz) and fast ripples (250-500 Hz) was calculated in MATLAB 2022b (MathWorks Inc., Natick, MA). The signal was segmented into continuous, non-overlapping 10-minute intervals. Each segment underwent further processing to determine the average power content within the designated frequency bands by employing the modified periodogram technique with a Hamming window. Additionally, alpha-delta ratio (ADR) and relative alpha variability (RAV) were assessed, with ADR quantifying the ratio of alpha to delta power within the selected EEG segment.

To quantify the number of fast ripples, data were filtered by removing noise using an automated MATLAB algorithm. Next, fast ripples were identified using the Short Time Energy (STE) method^25^. After identifying potential fast ripples, these detections were cross-checked using RIPPLELAB and then manually validated. Fast ripple analysis was performed using 24-hour continuous EEG recordings at three time points: 4 days post-injury in TBI mice (seizure-free), 4 days post-injury in seizure-experiencing mice (pre-seizure latent period) and at seizure onset in seizure-experiencing mice.

### Phase-Amplitude Coupling Analysis of Delta Waves and Fast Ripples

The timing of each fast ripple was analyzed to determine its coupling with delta waves. We extracted a 0.8 second window around each FR and filtered the data to isolate delta (1-4 Hz) and fast ripples (250-500 Hz) components. We assessed if fast ripples occurred during a peak-to-trough (DOWN state) or trough-to-peak (UP state) transition of delta using Hilbert transform at each fast ripple time.

### Statistical Analysis

To assess seizure occurrences during light or dark periods, we applied two-tailed Wilcoxon and binomial tests. We used one way ANOVA to compare spike counts among SHAM, TBI, and mice with seizures during the latent period. Spike counts between the latent period, and the seizure onset phase were compared with a two tailed t test. For fast ripples, we compared counts and their coupling to delta during up and down states across three groups: TBI, mice with seizures before onset (latent), and mice with seizures after onset using one way ANOVA.

## Results

### Characteristics of seizures after rdTBI

On day 0, mice underwent both rdTBI and EEG surgery. From day 1, we recorded video-EEG for up to 4½ months to track brain activity and seizures (**Fig. 1A**). EEGs used a monopolar setup with two recording electrodes plus ground and reference (**Fig. 1B**). **Fig. 1C** summarizes the distribution of animals across experimental groups. In total, 29 mice received rdTBI. Among these, 8 developed seizures during monitoring. 6 of those mice experienced recurrent seizures, and we labeled them as post-traumatic epilepsy (PTE). The remaining 2 mice had only a single seizure event and were labeled as post-traumatic seizure (PTS). 21 rdTBI mice, did not develop seizures. For comparison, 13 mice underwent sham procedures and served as controls (**Fig. 1C**). To characterize post-traumatic seizures after rdTBI, we measured the latency, onset, number, frequency, and incidence of seizures following established PTE classification criteria (**Fig. 1D**). Seizures were classified as early (≤7 days post-TBI) or late (>7 days post-TBI), with PTE defined as ≥2 unprovoked seizures occurring on separate occasions AND at least one seizure occurring >7 days after injury. Two mice experienced seizures within 7 days post-TBI. One mouse had a single early seizure (onset day 5) with no subsequent seizure activity during 115 days of monitoring and was classified as having an early single seizure. The second mouse had seizure onset on day 5 but developed 11 total seizures over 115 days of recording, meeting PTE criteria due to seizure recurrence. Five mice developed PTE with late seizure onset on days 8, 40, 80, 81, and 100, respectively, all exhibiting recurrent seizures during the monitoring period. One additional mouse experienced a single late seizure on day 126 with no subsequent seizure activity and was classified as having a single late seizure rather than PTE. Including the mouse with early seizure onset that progressed to recurrent seizures, the overall incidence of PTE was 20.6.% of injured mice (6/29 total TBI mice). For mice with late-onset PTE, seizures occurred with a mean latency period of 75.25 ± 12.7 days (mean ± SEM) from TBI. All PTE mice exhibited 6.8 ± 4.4 seizures (mean ± SEM) during their monitoring periods. **Fig. 1E** illustrates a representative seizure event recorded from one of the affected mice.

**Figure 1.**
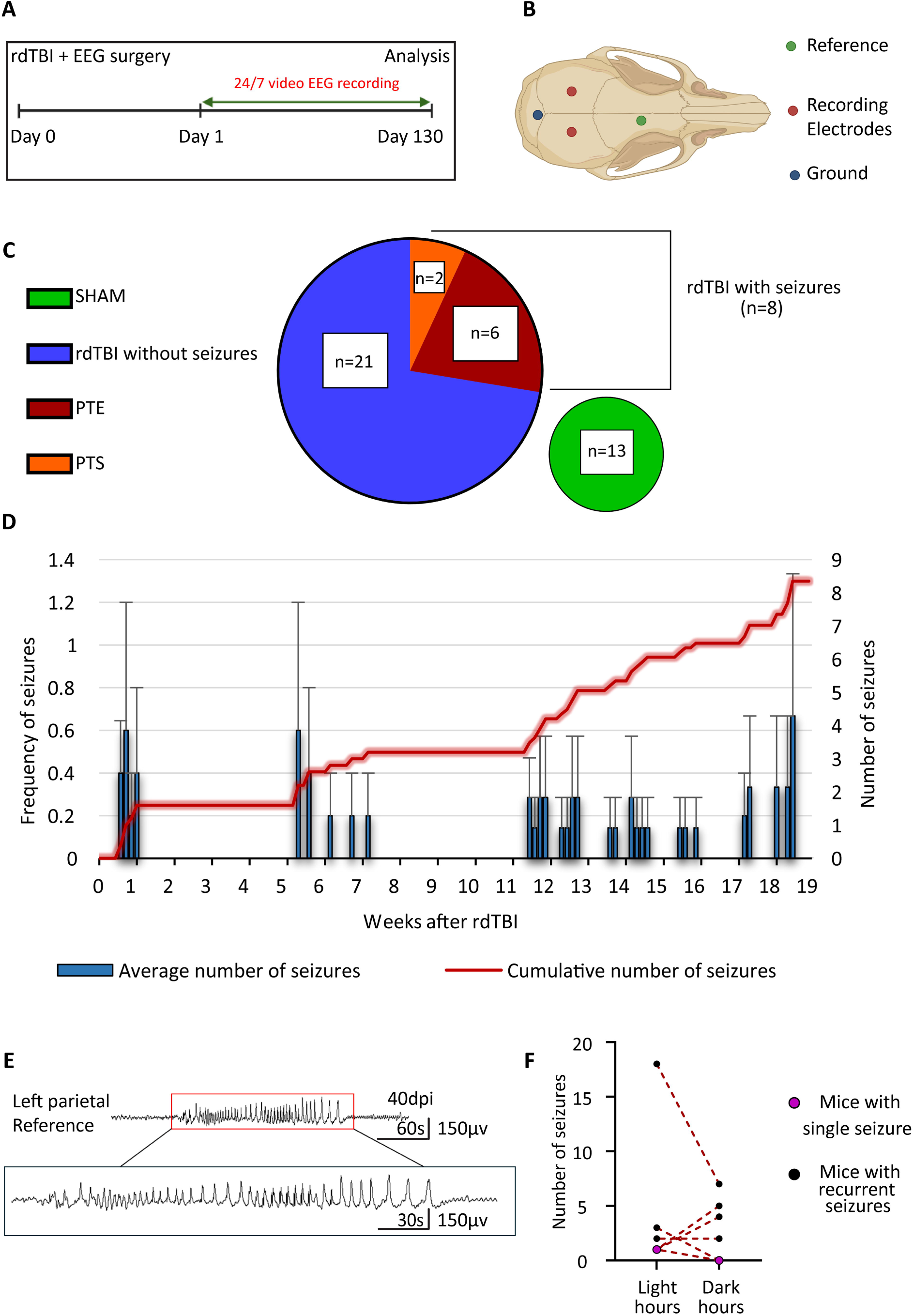
Seizure Incidence and Diurnal Patterns After rdTBI. (A) Mice underwent repetitive diffuse traumatic brain injury (rdTBI) and EEG surgery on the same day, followed by continuous 24/7 EEG-video monitoring for 130 days. (B) EEG electrode implantation was performed using monopolar montage with 2 recording electrodes (red), one ground (blue), and one reference (green) electrode. (C) In total, 29 mice underwent rdTBI. 8 developed seizures, with 6 progressing to post-traumatic epilepsy (PTE) and 2 showing only a single seizure, classified as post-traumatic seizure (PTS). The remaining 21 rdTBI mice did not develop seizures, and 13 additional mice served as sham controls. (D) Average number of seizures per mouse per week (mean ± SEM). (E) EEG trace displaying seizure in mice. (F) Comparison of seizure occurrences during the light (zeitgeber [ZT] 0-11) and dark (ZT 12-23) phases. No significant difference in median seizure numbers between light and dark phases was detected (two-tailed Wilcoxon test, p = 0.75).

### Diurnal seizure patterns after rdTBI

We next investigated seizures during different phases of the day. We found no significant difference in the seizure numbers’ median value between the light and dark phases (two-tailed Wilcoxon test, which does not *a priori* assume the direction of the difference, p = 0.75, **Fig. 1F**). Some mice had more seizures in the dark while others had more during the light phase. Yet, we observed that all mice experienced at least one seizure during the light phase and 3/8 (37.5%) mice had seizures exclusively during light hours, indicating a potential non-random pattern. We hypothesized that if there was no preference of seizure occurrence in either phase, then p will be 0.5 or the probability that an animal would experience one seizure at least in each phase is 50%. We determined that during the light phase the statistical probability of a seizure was significantly higher (two-tail binomial test, p = 0.0156). In contrast, no deviation from chance was determined for the dark phase (two-tail binomial test, p = 1.0). This suggests that individual mice are more likely to experience seizures during light hours.

### Seizure Onset Drives Further Spike Elevation in PTE

To characterize electrographic changes associated with PTE development, we analyzed spike activity patterns across experimental groups and disease progression phases. An example of typical epileptic spike in our mice is shown in **Fig. 2A**. One-way ANOVA revealed significant differences in 24-hour spike counts among experimental groups (Welch’s ANOVA: W (2, 9.913) = 15.45, p = 0.0009). Post-hoc analysis using Dunnett’s T3 multiple comparisons test demonstrated that TBI groups exhibited significantly elevated spike activity compared to SHAM controls (**Fig. 2B**). SHAM mice showed minimal baseline spike activity (1.0 ± 0.2 spikes per 24 hours, n = 4). TBI mice that did not develop PTE exhibited a significant 5.4-fold increase in spike frequency compared to SHAM (6.4 ± 0.8 spikes, mean difference = −5.364, 95% CI [-9.185 to −1.542], p = 0.0067), indicating that traumatic brain injury itself induces cortical hyperexcitability independent of seizure development. PTE mice during the pre-onset latent phase showed the highest spike activity (8.5 ± 1.0 spikes, n = 6), representing a significant 7.5-fold increase compared to SHAM controls (mean difference = −7.500, 95% CI [-12.79 to −2.206], p = 0.0110). However, the difference between TBI and PTE pre-onset groups did not reach statistical significance (mean difference = −2.136, 95% CI [-7.944 to 3.671], p = 0.6753), suggesting that spike frequency alone during the latent period cannot reliably distinguish mice that will develop PTE from those that will not.

**Figure 2.**
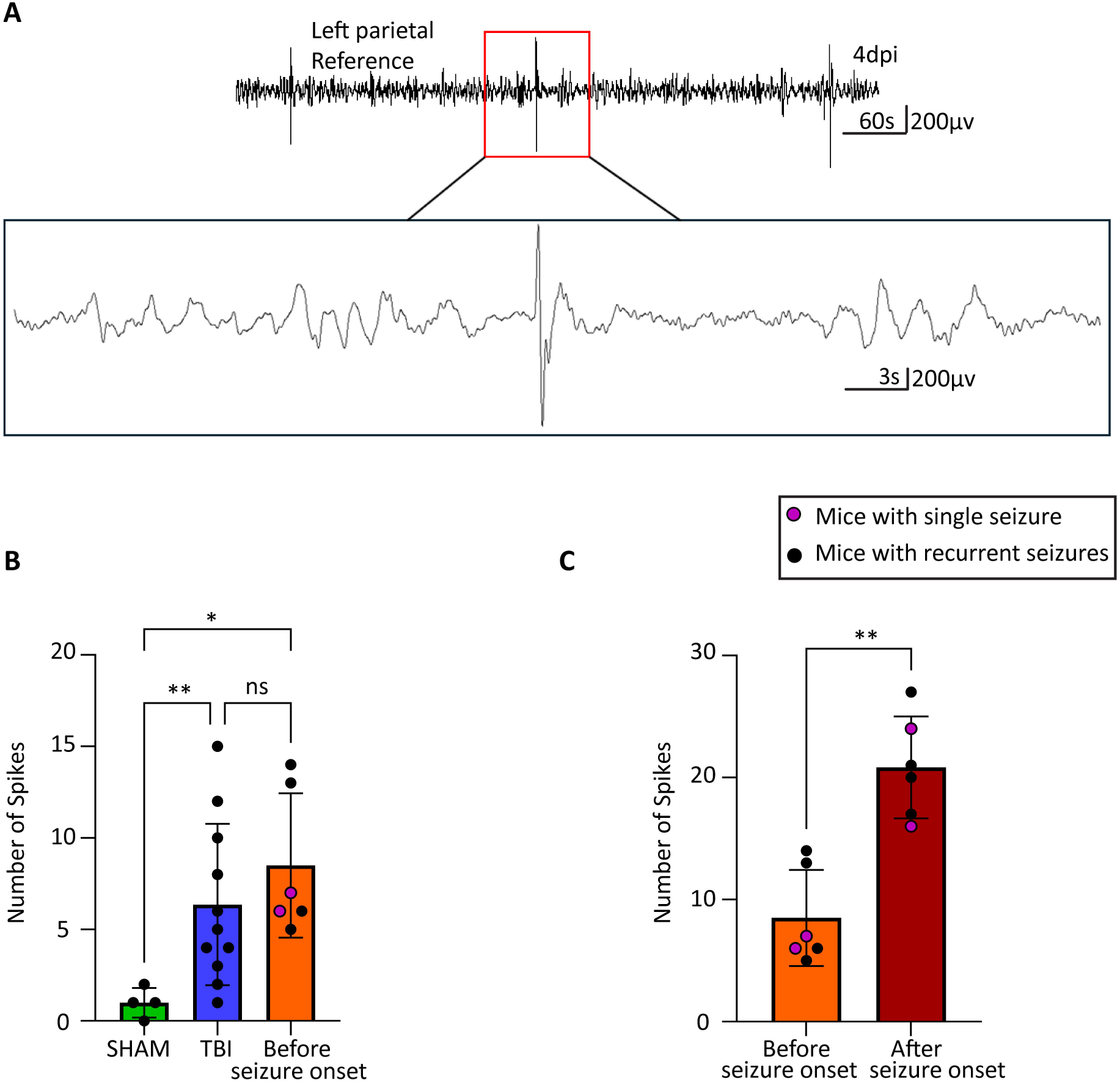
Spike comparison across animal groups. (A) Representative trace of typical epileptic spikes in PTE mice. (B) At 4 days post injury (dpi), SHAM mice had significantly fewer spikes than TBI mice (one-way ANOVA, p = 0.0067; n = 4 SHAM, n = 11 TBI) and mice with seizures in the latent phase (one-way ANOVA, p = 0.0110; n = 4 SHAM, n = 6 latent). No difference was found between TBI mice and latent phase mice (one-way ANOVA, p = 0.6753; n = 11 TBI, n = 6 latent). (C) The number of spikes was significantly higher in mice with seizures after seizure onset compared to the latent phase (two-tailed t-test, t=5.940, df=5, p=0.0019, n=6 per group).

A more pronounced change occurred when comparing spike activity before and after seizure onset within the same PTE mice (**Fig. 2C**). A paired t-test comparing spike counts in PTE mice pre- and post-seizure onset revealed a significant increase in spike frequency (mean increase = +12.33 spikes; 95% CI [6.996, 17.67]; t₅ = 5.94, p = 0.0019, η²ₚ = 0.876). This represents an exceptionally large effect size (Cohen’s d = 2.43), with seizure onset accounting for approximately 88% of the variance in spike frequency changes. Spike activity increased from 7.3 ± 1.2 spikes during the pre-onset phase to 20.2 ± 1.7 spikes following seizure development, representing a nearly 3-fold escalation in interictal hyperexcitability. This dramatic increase suggests that the emergence of spontaneous seizures triggers a cascade of network instability that extends well beyond ictal periods.

These findings reveal a biphasic pattern: initial TBI-induced spike elevation affecting all injured animals, followed by dramatic secondary escalation after seizure onset in PTE mice. This post-seizure increase represents establishment of chronic epileptic networks, supporting epileptogenesis as a dynamic process where seizures facilitate further network destabilization. Yet, given the lack of significant difference between TBI and PTE mice during the pre-seizure latent period (p = 0.6753), spike counts alone are insufficient as predictive biomarkers for PTE development.

### EEG Biomarker Analysis in PTE Through Power Spectrum Changes

For EEG biomarker analysis, we aimed to determine whether power spectrum differences could distinguish mice that developed post-traumatic seizures (regardless of seizure pattern - single or recurrent) from TBI mice that remained seizure-free throughout the observation period. We analyzed EEG power spectrum changes across multiple frequency bands over 17 weeks using an automated MATLAB algorithm to identify potential electrographic signatures that precede or accompany seizure development (**Fig. 3**). EEG recordings from SHAM, TBI and seizure-experiencing groups were analyzed across delta (0.5-4 Hz), theta (4.5-7.5 Hz), alpha (8-13 Hz), beta (13-30 Hz), gamma (30-80 Hz) and high-frequency oscillations (HFO; 100-200 Hz) bands, along with derived metrics including alpha-delta ratio (ADR), relative alpha variability (RAV) and total power.

**Figure 3.**
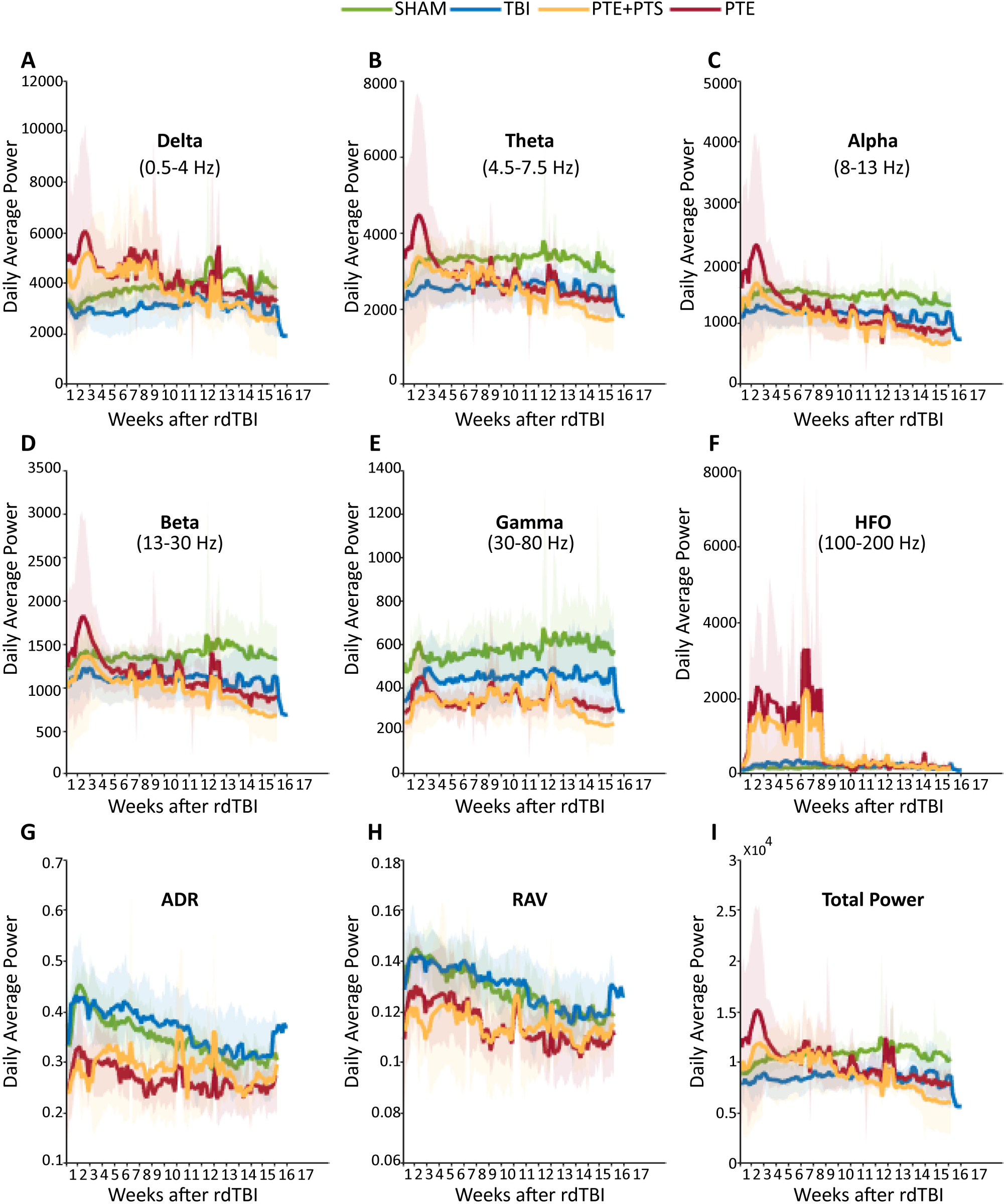
EEG Power Spectrum Trends in PTE mice. Average power changes over the 17-week period post-rdTBI for (A) delta (0.5 - 4 Hz), (B) theta (4.5 - 7.5 Hz), (C) alpha (8 - 13 Hz), (D) beta (13 - 30 Hz), (E) gamma (30 - 80 Hz), (F) high-frequency oscillations (HFO; 100 - 200 Hz), (G) ADR (Alpha-Delta Ratio), (H) RAV (Relative Alpha Variability), and (I) total power. Power is presented as mean (lines) with standard deviation (cloud) for each group: SHAM (n=4, green), TBI (n=11, blue), PTE+PTS (n=6, orange) and PTE (n=4, red). When the two post traumatic seizure (PTS) mice were excluded from the post traumatic epilepsy (PTE) group, the overall trend remained unchanged, suggesting PTS and PTE share similar features.

Seizure-experiencing mice exhibited distinctly elevated delta power compared to both SHAM and seizure-free TBI mice, particularly during early post-injury weeks (**Fig. 3A**). This slow-wave elevation was sustained throughout monitoring, with episodic spikes around weeks 8 and 12 likely corresponding to increased seizure activity. Similarly, theta power showed consistent elevations in seizure-experiencing mice compared to seizure-free TBI mice (**Fig. 3B**), suggesting slow-wave activity as an early electrographic marker of seizure susceptibility. Conversely, seizure-experiencing mice demonstrated consistently reduced power in faster physiological frequency bands. Alpha, beta and gamma powers were all diminished in seizure-experiencing mice relative to both SHAM and seizure-free TBI groups (**Fig. 3C, D, E**), suggesting progressive cortical network dysfunction in seizure-prone mice. Paradoxically, while physiological gamma was reduced, HFO were markedly elevated in seizure-experiencing mice, with prominent peaks around weeks 2, 4 and 6 (**Fig. 3F**). This shift toward pathological HFO activity, combined with reduced physiological gamma, indicates fundamental network excitability alterations distinguishing seizure-prone from seizure-resistant TBI mice. ADR and RAV both showed sustained reductions in seizure-experiencing mice compared to controls (**Fig. 3G, H**), effectively capturing the dual pattern characterizing mice destined for post-traumatic seizures. Total power showed decreased trends in seizure-experiencing mice compared to TBI and SHAM (**Fig. 3I**).

### Fast Ripple Dynamics and Delta Coupling in Post-Traumatic Epilepsy

Fast ripples (250-500 Hz) are pathological HFO distinct from physiological HFO and interictal spikes, representing synchronized activity of small neuronal populations that has been associated with epileptogenic tissue in both human patients^18, 26^ and animal models^27^. Unlike spikes, which are brief transient discharges, fast ripples are sustained oscillatory events that may reflect underlying network hyperexcitability. A representative trace of fast ripples in RIPPLELAB has been shown in **Fig. 4A**. We quantified fast ripples at three time points: 4 days post-TBI (latent period) and at seizure onset in seizure-experiencing mice. Fast ripple frequency was significantly elevated in seizure-experiencing mice compared to seizure-free TBI mice both during the latent period (**Fig. 4B**, p=0.0018) and at seizure onset (**Fig. 4B**, p=0.0038), establishing fast ripples as a consistent biomarker across the epileptogenic process. To examine the temporal relationship between fast ripples and slow-wave activity, we analyzed phase-amplitude coupling between fast ripples and delta waves (1-4 Hz). Delta oscillations alternate between UP states (depolarized, high-amplitude) and DOWN states (hyperpolarized, low-amplitude), representing different phases of cortical excitability. Examples of coupling in TBI and PTE mice have been shown in **Fig. 4C**. While fast ripple coupling to delta UP states showed no significant differences between groups (**Fig. 4D**), fast ripple coupling to delta DOWN states was significantly increased in seizure-experiencing mice both during the latent period (**Fig. 4E**, p=0.0082) and after seizure onset (**Fig. 4E**, p=0.0392) compared to seizure-free TBI mice.

**Figure 4:**
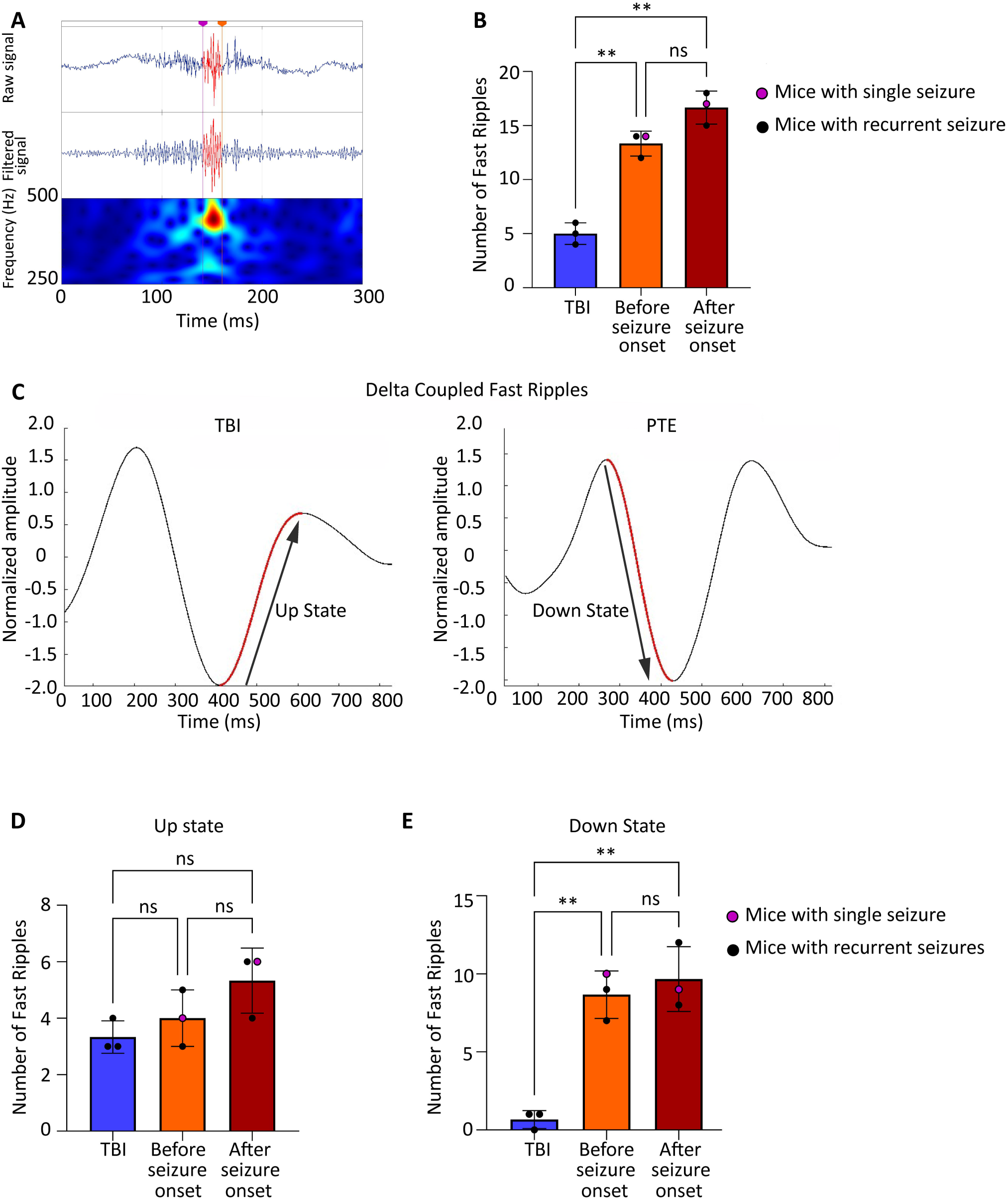
Fast ripple dynamics and delta coupling in PTE. (A) Representative EEG trace showing fast ripple events in PTE mice. (B) Fast ripples were significantly higher in seizure experiencing mice compared to TBI during the latent phase (p = 0.0018, n = 3 per group) and remained elevated after seizure onset (p = 0.0038), with no difference between latent and post onset phases (p = 0.0963) which suggests sustained fast ripples activity during both periods. (C) Normalized delta amplitude waveforms coupled with fast ripples, showing differences between TBI and PTE mice. (D) Fast ripples coupling to delta UP states showed no significant difference between seizure and TBI mice during the latent phase (p = 0.7093) or after seizure onset (p = 0.1686). (E) Fast ripples coupling to delta DOWN states were significantly increased in seizure mice during the latent phase (p = 0.0082) and remained higher after onset (*p* = 0.0390), with no difference between pre and post onset phases (p = 0.8733).

This preferential coupling of fast ripples to delta DOWN states suggests that pathological high-frequency activity emerges during periods of cortical inhibition, potentially reflecting a breakdown in normal inhibitory control mechanisms. The presence of this coupling signature during the latent period, before any seizures occur, establishes fast ripple-delta DOWN state coupling as a promising early biomarker for post-traumatic seizure susceptibility.

## Discussion

Here, we assessed differences in EEG signatures between TBI animals that do or do not develop seizures for their usefulness as early biomarkers. Our findings, in concert with those of recent investigations in focal epilepsy^18^, indicate that fast ripples, especially delta-coupled fast ripples, may be early biomarkers for epileptogenesis. In human patients with epilepsy, HFOs, especially fast ripples, were suggested as biomarkers of the seizure onset zone^28^. They were also recorded in various rodent models of epilepsy^18^.

Delta coupling analysis revealed distinct patterns between TBI and seizure-experiencing mice, with a preference for coupling to the delta DOWN state rather than the UP state before seizure onset. This is particularly interesting because delta DOWN-state coupling of fast ripples emerged before seizure onset, suggesting early network instability specific to the latent phase of post-traumatic seizure development. After seizure onset, coupling shifted further toward the delta DOWN state, indicating a dynamic interaction between these oscillations and the progression of epileptogenesis. Disruptions in delta wave coupling may reflect changes in thalamocortical network dynamics that may contribute to the persistence of hyperexcitability. This aligns with prior reports that these states are critical for initiating epileptiform activity, as they represent periods of synchronized cortical inhibition that can facilitate seizure propagation^29^.

We found significant elevations in delta power during the early post-injury weeks in seizure-experiencing animals, supporting its role as a biomarker for epileptogenesis^30^. The sustained increase in delta activity provides evidence of its association with chronic network dysfunction^31^. Additionally, the decreased power observed in alpha and beta frequency bands aligns with findings of cortical network disruptions that are frequently observed in post-TBI studies^32^.

Gamma-to-HFO ratio reductions and consistent gamma band deficits may reflect disrupted GABAergic inhibition within cortical networks, as gamma oscillations are typically associated with inhibitory interneuron activity^33^. A reduction in gamma power alongside increased HFO suggests an imbalance between excitatory and inhibitory signaling, potentially driven by a loss of GABA-mediated control, leading to increased network excitability and seizure susceptibility. These findings extend the predictive capability of EEG analysis by highlighting shifts in physiological oscillations toward pathological activity, an observation also reported in human qEEG studies of post-traumatic seizure disorders^31^.

Critically, our power spectrum analysis demonstrated that mice experiencing any post-traumatic seizure, whether single early, single late or recurrent, exhibited similar patterns of brain network dysfunction compared to seizure-free TBI animals. These seizure-experiencing mice showed elevated delta and theta power, reduced physiological fast oscillations and increased pathological HFO activity, regardless of seizure frequency or timing. This electrographic similarity challenges the assumption that single seizures represent fundamentally different pathophysiological processes from recurrent seizures.

Our spike analysis exemplified the limitation of individual biomarkers: while seizure-experiencing mice showed dramatic increases in spike activity after seizure onset, the overall difference between TBI (seizure-free) and pre-seizure PTE groups was non-significant, suggesting that spikes count alone during the latent period cannot reliably distinguish which animals will develop seizures from those that will not. This highlights a fundamental challenge in PTE biomarker development, individual electrographic measures may lack sufficient sensitivity during the critical pre-seizure window when intervention would be most valuable. The fast ripple-delta coupling we identified as an early biomarker emerged precisely because we examined more sophisticated temporal relationships rather than relying on simple power measurements or event counts during arbitrary time windows.

While our study included animals with single seizures (n=2), precluding statistical comparison, both exhibited similar electrographic abnormalities (elevated fast ripples and delta-coupling) as PTE mice. This suggests single and recurrent post-traumatic seizures may represent the same pathophysiological spectrum, warranting re-examination of current binary classification systems. The considerable heterogeneity we observed (seizure onset: 5-126 days post-injury) reflects post-traumatic epileptogenesis complexity seen in patients, indicating biomarker strategies must account for individual trajectories rather than uniform pathophysiology.

Our current findings build upon our previous research^9, 10^, where continuous EEG monitoring was limited to a two-month period post-injury. The extension of monitoring up to 19 weeks in the present study has allowed for a more comprehensive analysis of fast ripples during the latent and chronic phases of seizure development. The variability in seizure onset times we observed aligns with the variability seen in human TBI cases^34–39^, highlighting the challenge of using rigid diagnostic criteria for these heterogeneous disorders.

In conclusion, we identified several EEG biomarkers that distinguished rdTBI mice experiencing any form of post-traumatic seizure from those that remained seizure-free, with fast ripple-delta coupling emerging as the most promising early biomarker. Our findings demonstrate that this biomarker can identify seizure-prone animals during the critical latent period when intervention would be most beneficial. Future research should validate these findings in larger cohorts with adequate sample sizes to determine whether single and recurrent post-traumatic seizures represent similar pathophysiological processes and whether the binary recurrence threshold for PTE diagnosis adequately captures the biological reality of post-traumatic network dysfunction.

## Abbreviations

ADR: Alpha-Delta Ratio
CDMRP: Congressionally Directed Medical Research Programs
CCI: Controlled Cortical Impact
CURE: Citizens United for Research in Epilepsy
DoD: Department of Defense
EEG: Electroencephalography
FPI: Fluid Percussion Injury
HFO: High-Frequency Oscillations
IIR: Infinite Impulse Response
NIH: National Institutes of Health
NINDS: National Institute of Neurological Disorders and Stroke
PTE: Post-Traumatic Epilepsy
PTS: Post-Traumatic Seizure
RAV: Relative Alpha Variability
rdTBI: Repetitive Diffuse Traumatic Brain Injury
SEM: Standard Error of the Mean
STE: Short Time Energy
TBI: Traumatic Brain Injury
ZT: Zeitgeber Time

## Acknowledgements

This work was supported by CURE Epilepsy based on a grant CURE Epilepsy received from the United States Army Medical Research and Materiel Command, Department of Defense (DoD), through the Psychological Health and Traumatic Brain Injury Research Program under Award No. W81XWH-15-2-0069. The work in this manuscript was further supported by the National Institutes of Health grant from the National Institute of Neurological Disorders and Stroke R01NS121145 to SR and the Department of Defense CDMRP #HT94252410116 to OS. Opinions, interpretations, conclusions and recommendations are those of the author and are not necessarily endorsed by the Department of Defense or the National Institutes of Health. In conducting research using animals, the investigator(s) adheres to the laws of the United States and regulations of the Department of Agriculture. OL was supported by the Cell and Molecular Biology Training Grant T32GM146611 at the University of Alabama at Birmingham.

## Conflict of Interest Statement

None.

## Data Availability Statement

The data that support the findings of this study are available from the corresponding author upon reasonable request.

## Notes

### Competing Interest Statement

The authors have declared no competing interest.

### Summary of Updates

This is a shorter version of the earlier paper, prepared to fit the word limit for Epilepsia Open.

